# Water Diffusion in the Live Human Brain is Gaussian at the Mesoscale

**DOI:** 10.1101/2024.04.10.588939

**Authors:** Kulam Najmudeen Magdoom, Alexandru V. Avram, Thomas E. Witzel, Susie Y. Huang, Peter J. Basser

**Affiliations:** Section on Quantitative Imaging and Tissue Sciences, The Eunice Kennedy Shriver National Institute of Child Health and Human Development, National Institutes of Health, Bethesda, MD, 20892 USA; The Military Traumatic Brain Injury Initiative (MTBI), Uniformed Services University of the Health Sciences, Bethesda, MD 20814, USA; The Henry M. Jackson Foundation for the Advancement of Military Medicine (HJF) Inc., Bethesda, MD, USA; Athinoula A. Martinos Center for Biomedical Imaging, Department of Radiology, Massachusetts General Hospital, Boston, MA, USA

**Keywords:** diffusion, Gaussian, mesoscale, MRI, pulsed-field gradient, mPFG, MDE, b-tensor, DTI, DTD, heterogeneity, anisotropy, DWI, microFA, MD distribution, b-matrix, b-tensor

## Abstract

Imaging the live human brain at the mesoscopic scale is a desideratum in basic and clinical neurosciences. Despite the promise of diffusion MRI, the lack of an accurate model relating the measured signal and the associated microstructure has hampered its success. The widely used diffusion tensor MRI (DTI) model assumes an anisotropic Gaussian diffusion process in each voxel, but lacks the ability to capture intravoxel heterogeneity. This study explores the extension of the DTI model to mesoscopic length scales by use of the diffusion tensor distribution (DTD) model, which assumes a Gaussian diffusion process in each subvoxel. DTD MRI has shown promise in addressing some limitations of DTI, particularly in distinguishing among different types of brain cancers and elucidating multiple fiber populations within a voxel. However, its validity in live brain tissue has never been established. Here, multiple diffusion-encoded (MDE) data were acquired in the living human brain using a 3 Tesla MRI scanner with large diffusion weighting factors. Two different diffusion times (Δ = 37, 74 ms) were employed, with other scanning parameters fixed to assess signal decay differences. In vivo diffusion-weighted signals in gray and white matter were nearly identical at the two diffusion times. Fitting the signals to the DTD model yielded indistinguishable results, except in the cerebrospinal fluid (CSF)-filled voxels likely due to pulsatile flow. Overall, the study supports the time invariance of water diffusion at the mesoscopic scale in live brain parenchyma, extending the validity of the anisotropic Gaussian diffusion model in clinical brain imaging.

## 1 Introduction

Imaging neural tissue microstructure *in vivo* is a holy grail for neuroscience and neuroradiology. Accessing neural tissue structure at the mesoscopic scale would enable the study of fundamental structure-function relationships in the living human brain and enable the development of early markers sensitive to a range of neuropathology. The diffusion of water molecules serves as a sensitive and non-invasive probe for tissue microstructure given the abundance of water within tissue and the microscopic nature of diffusion displacements. A handful of methods currently exist to measure diffusive displacements of indicator molecules in biological tissues. Fluorescent methods [1, 2] provide the specificity to follow single particles but often require exogenous dyes and operate in a limited field of view (FOV). MRI methods using pulsed-field gradients (PFG) do not have the specificity of fluorescent methods but are scalable to the whole human brain, use tissue water as an intrinsic contrast agent, and are well-suited to study intermediate mesoscopic brain architecture and circuits *in vivo* [3]. However, the lack of specificity of MRI methods makes the reconstruction of the underlying tissue microstructure from measured diffusion displacements ill-posed, akin to the problem of ‘hearing the shape of a drum’ [4]. An understanding of the nature of water diffusion in brain tissue *in vivo* at mesoscopic length and time scales is critical to solving this important problem.

It is well known that the diffusivity of water in brain tissue is less than that in free water, thus indicating the presence of diffusive barriers [5], which are largely attributed to lipid bilayers that envelope a myriad of microscopic compartments observed in electron micrographs of brain tissue specimens. Evidence for this comes from experiments in which lipids are removed from tissue using surfactants, after which the value of tissue diffusivity is close to that of free water [6, 7]. The diffusivity of water in brain parenchyma is also anisotropic as evidenced by an orientation-dependent diffusion-weighted MR signal [8]. Diffusion tensor imaging (DTI), which assumed Gaussian diffusion displacements of water molecules [9, 10] was the first comprehensive model to quantify this diffusion anisotropy in living brain tissue, especially in white matter [11]. The inability of the DTI model to disambiguate certain tissue microstructural motifs led to the development of several models [12, 13, 14] that questioned the core assumption of the DTI model, *i.e.*, the presence of Gaussian diffusion compartments in living brain tissue.

A signature of Gaussian diffusion is that the diffusion process satisfies the Einstein formula, a linear relationship between the mean-squared net displacement and the diffusion time (*i.e.*, resulting in a constant diffusivity). MRI studies measuring the time dependence of water diffusivity in brain tissue have been inconsistent, likely due to differences in the length and time scales probed, data analysis methods used, sample type (fixed vs. live tissue), and MRI scanning parameters. These studies, however, demonstrate the general trend that the apparent water diffusivity in the brain decreases with diffusion time at very short diffusion times and plateaus to a constant non-zero value thereafter [15, 16, 17, 18]. Several models have been proposed to explain the observed trend [13, 19, 20]. Nevertheless, it is clear that water diffusion in the brain may not be fully restricted as in glass capillary arrays [21, 22], since such impermeable barriers would result in a continuous reduction in diffusivity with respect to the diffusion time, approaching zero at very long diffusion times.

Given the constancy of tissue water diffusivity over a wide range of time scales, the central question we address in this work is whether a Gaussian diffusion assumption in brain tissues is justified at the mesoscale. We address this question using an advanced human MRI scanner optimized for acquiring data with large diffusion weighting factors with a high signal-to-noise ratio (SNR) (*i.e.*, the Connectome MRI scanner [23]). The advantage of the Gaussian diffusion model is its simplicity and ability to capture anisotropy known to exist in neural tissue [9, 24]. The limitations of the Gaussian diffusion-based DTI model are largely attributed to the assumption of structural homogeneity within a voxel [25, 26], and the inability of single-PFG signals to distinguish between and among different underlying microstructural motifs [27, 28, 29].

Jian et al. [30] introduced a new model to overcome these limitations of DTI by invoking Gaussian diffusion within microvoxels that constitute the macroscopic MRI voxel which is described by a probability density function of diffusion tensors (i.e., DTD [31]). Despite its introduction two decades ago and its ability to distinguish different types of brain tumors using multiple PFG measurements [32], the validity of this model for brain tissue especially the use of exponential kernel to describe the signal has not been investigated. If diffusion were Gaussian within mesoscopic “microvoxels” that constitute a larger imaging voxel, one would predict that each of these microscopic compartments would exhibit time-independent diffusion, and thus their aggregate or ensemble-averaged DTD would also be independent of diffusion time, which is the hypothesis we test in this paper. We acquired diffusion-weighted (DW) MR signals in the living human brain at two widely different diffusion times and using strong diffusion weighting factors (*i.e.*, b-value) using single- and double-PFG MRI acquisitions. These DW signals were then used to estimate the DTD, which we assume to be the maximum entropy normal tensor-variate distribution (NTVD) with samples constrained to be positive definite (CNTVD) [33]. This assumption ensures physicality since negative-definite diffusion tensors are inadmissible as they would violate the second law of thermodynamics.

## 2 Materials and Methods

### 2.1 DTD signal model

The MR signal from the probability density of independent components of the diffusion tensor, *p* (D), is given by [30],

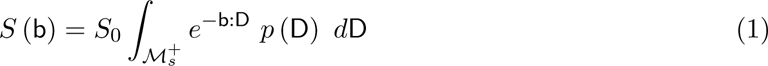

where *S*_0_ is the signal without diffusion weighting, 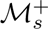 is the space of symmetric positive definite diffusion tensors, and b : D is the tensor dot product of the symmetric second-order b-tensor and diffusion tensor, respectively. Assuming a CNTVD for *p* (D), the signal equation is approximated using Monte Carlo (MC) integration with samples, D, drawn from the NTVD [31] with a given second-order mean, D̅, and fourth-order covariance tensor, Ω, which are filtered as shown below to ensure positive definiteness of the sample diffusion tensors comprising the ensemble [29]. In other words, the diffusion-weighted signal is the expected value of the exponential Gaussian diffusion kernel whose tensor random variables are drawn from a normal tensor variate distribution that is sampled over a domain of positive definite tensors,

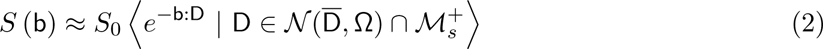

where the angular brackets denote ensemble averaging over all micro-diffusion tensors, and *N* is the NTVD. For convenience, the mean and covariance tensors of *N* are expressed as a 6D vector and 6 *×* 6 symmetric positive definite matrix, respectively, when drawing samples from the NTVD. It should be noted that time-dependent diffusion in the brain would render the exponential Gaussian diffusion kernel invalid, which is the subject of our investigation.

### 2.2 MRI measurements and pre-processing

MRI data was acquired using a 64-channel coil on the 3T Connectome MRI system (MAGNETOM Connectom, Siemens Healthineers) that produces a maximum 300 mT/m gradient strength and 200 T/m/s slew rate. Diffusion MRI was performed on five healthy young adults (21-24 years of age, 4 females) who provided written informed consent in accordance with a research protocol approved by the Institutional Review Board (IRB) of Mass General Brigham. The human DWI data was anonymized prior to receipt and subsequent analysis.

Diffusion MRI acquisition was also performed on 40% polyvinylpyrrolidone phantom at ambient bore temperature for control purposes [34]. The DTD MRI acquisition was performed using interfused-PFG (iPFG) [33] incorporated into a multi-band slice selective echo-planar imaging (MB-EPI) sequence [35] for efficient whole brain coverage. The well-defined diffusion time in iPFG allowed us to perform the DTD MRI acquisition at two different diffusion times while fixing all other scan parameters as shown in Figure 1.

**Figure 1:**
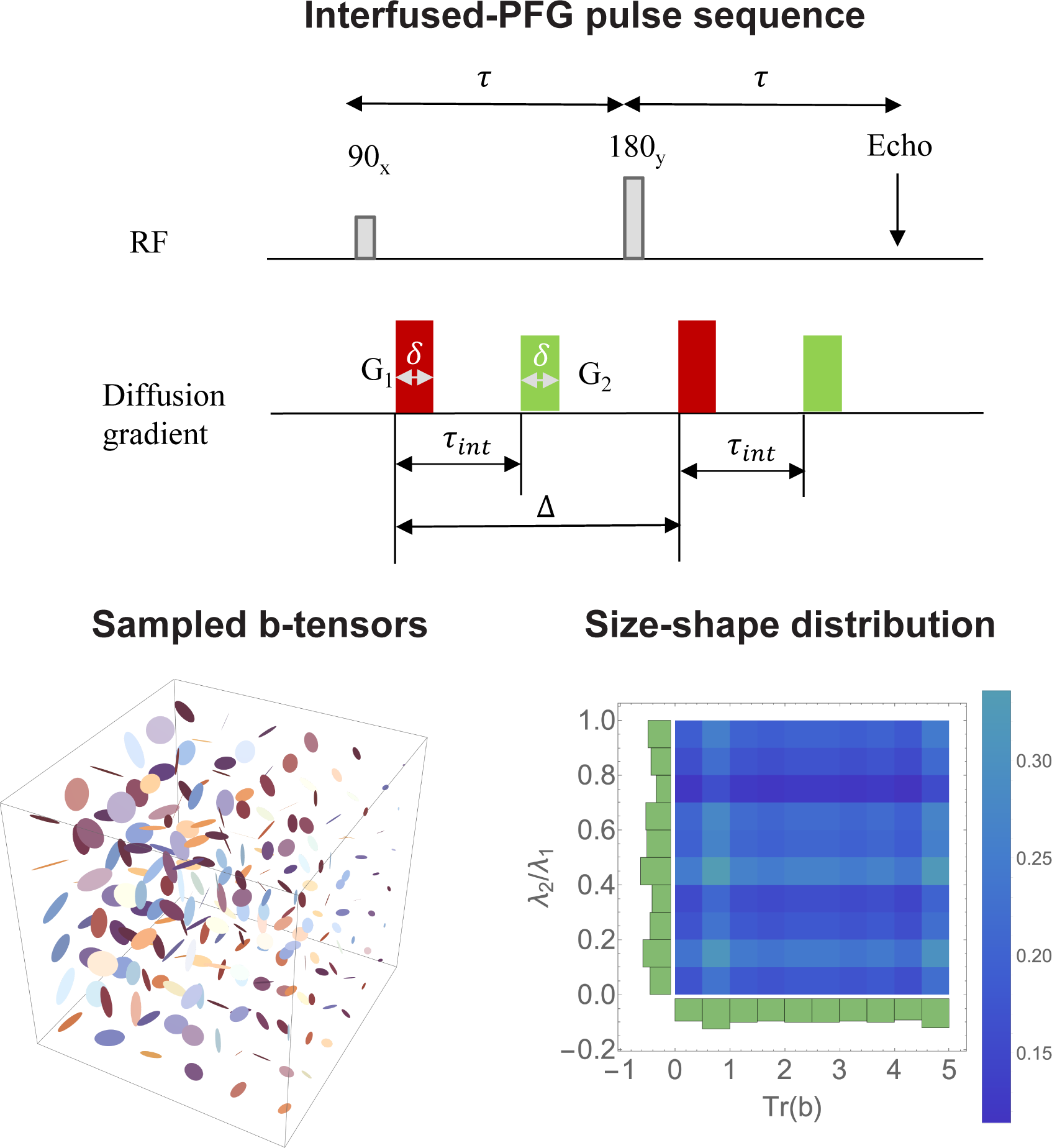
Interfused pulsed-field gradient (iPFG) sequence used to realize the rank-1 and rank-2 b-tensors that are displayed as sticks and ellipses, respectively. The well-defined diffusion timing parameters in the sequence let us realize the same b-tensor at different diffusion times while fixing the remaining parameters. The parameters, *G*_1_ and *G*_2_ are the two independent diffusion gradient magnitudes, *τ*_int_ is the time between the two gradients (i.e., interfusing time), *δ* is the diffusion gradient duration, and Δ is the diffusion time. The uniform size-shape distribution of the b-tensors sampled at both diffusion times is displayed as a 2D histogram of trace(b) and the ratio of non-zero eigenvalues of the b-tensor.

A total of 216 axial diffusion-weighted images (DWIs) were acquired with rank-1 and rank-2 b-tensors of uniform size, shape, and orientation with b-values ranging from 0 – 5,000 *s/mm*^2^ (Figure 1) using the following parameters: *δ* = 11 ms, Δ = 37, 74 ms, FOV = 192 *×* 192 *×* 140 mm, MB factor = 3, TR*\*TE = 4,600*\*145 ms at 2 mm isotropic spatial resolution. For comparison, all the DWIs for a given subject at both the diffusion times were registered with the non-diffusion weighted imaging volume acquired at Δ = 37 ms using the 3D affine registration tool [36, 37] available in FSL software environment [38].

### 2.3 Microstructural measures derived from the DTD

The mean and covariance tensors characterizing the CNTVD at the two diffusion times were estimated from the DWIs as outlined in [33]. The estimated CNTVD parameters were used to delineate several microstructural features within each voxel. The micro-diffusion tensors in the voxel were simulated by drawing MC samples from the CNTVD with the estimated mean and covariance tensors. The tissue heterogeneity was classified based on the size, shape, and orientation heterogeneity of the micro-diffusion tensor ellipsoids. The size heterogeneity of micro-diffusion tensors was quantified by the mean and standard deviation of one-third of the trace (*i.e.*, mean diffusivity, MD, and its standard deviation, *σ*_MD_). The shape and orientation heterogeneity was quantified by the microscopic fractional anisotropy (*µ*FA) which was computed directly from the ensemble of individual fractional anisotropies (FA), *i.e.*, averaged over each of the micro-diffusion tensors. The “macro” and “micro” FA are generally not equal due to the non-commutativity of the FA operator.

## 3 Results

In this section, we present the raw DWIs, and DTI (*i.e.*, MD and FA)/DTD-MRI derived parameters (*i.e.*, *σ*_MD_ and *µ*FA) from a representative axial slice acquired at the two diffusion times in our PVP phantom and *in-vivo* human brains. A representative set of four b-tensors was chosen for raw DWIs which includes both rank-1 and rank-2 with small, medium, and large b-values. The absolute difference between the DWIs acquired at both diffusion times and the distribution of their intensities are also shown to delineate the differences in the signal between the two diffusion times. The DTI and DTD measures were also quantitatively compared across the diffusion times by reporting their values averaged over a region of interest (ROI). Finally, the variability in the results across the population was demonstrated by showing the DWI difference maps across the five subjects scanned for the four b-tensors chosen previously.

The raw DWIs obtained in the macroscopically and microscopically isotropic PVP phantom for the various b-tensors are shown in Figure 2. The images from the two diffusion times were nearly identical as shown by the absolute difference maps in the third row in the figure. There were small differences (*<* 3%) observed at the lower b-values which were manifest in the shifts observed in the histogram of the signal differences. There were also some differences at the boundaries that may be attributed to imperfect image registration. The DTI and DTD measures were nearly identical for both diffusion times as shown by the maps and ROI averaged values in the center of the phantom shown in Figure 3 and Table 1 respectively. The value of the mean diffusivity was 0.58 *±* 0.01*µm*^2^*/ms*, while the FA and *µ*FA was approximately 0.03 and 0.04 respectively. Image ghosting was observed on *S*_0_ and mean diffusivity spectral moments.

**Figure 2:**
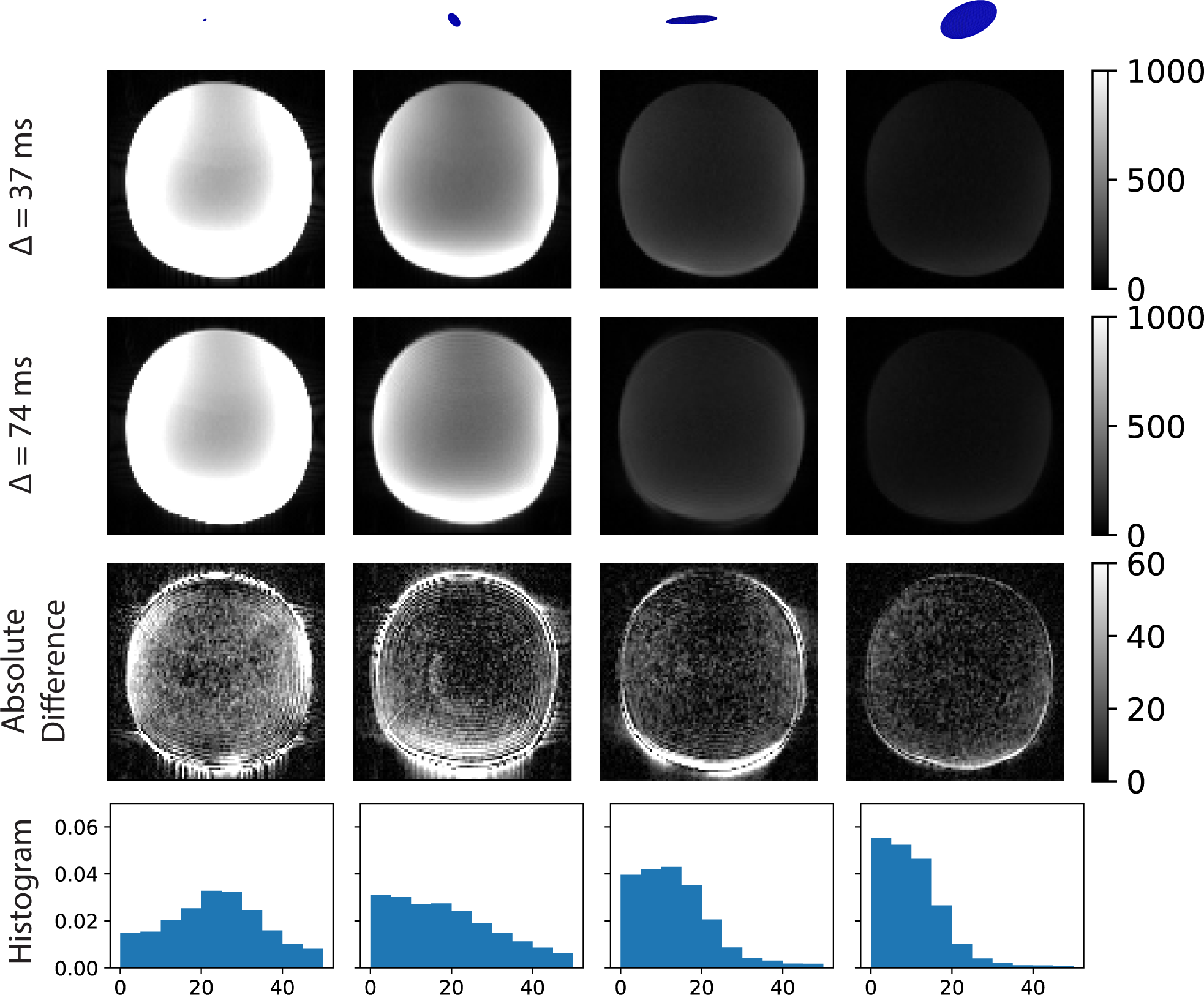
Multiple diffusion encoded (MDE) images of an axial slice of a PVP phantom acquired at two different diffusion times (i.e., Δ = 37*\*74 ms). The rank-1/rank-2 b-tensors corresponding to each column are displayed as ellipses. The b-values of the plotted b-tensors are approximately 270, 1100, 3500, and 4820 *s/mm*^2^ respectively. The absolute difference between the two images along with its histogram are shown in the third and fourth rows respectively. DWIs at the two diffusion times for the acquired b-tensors were indistinguishable as shown in the absolute difference image except at the boundaries which might be due to imperfect image registration.

**Figure 3:**
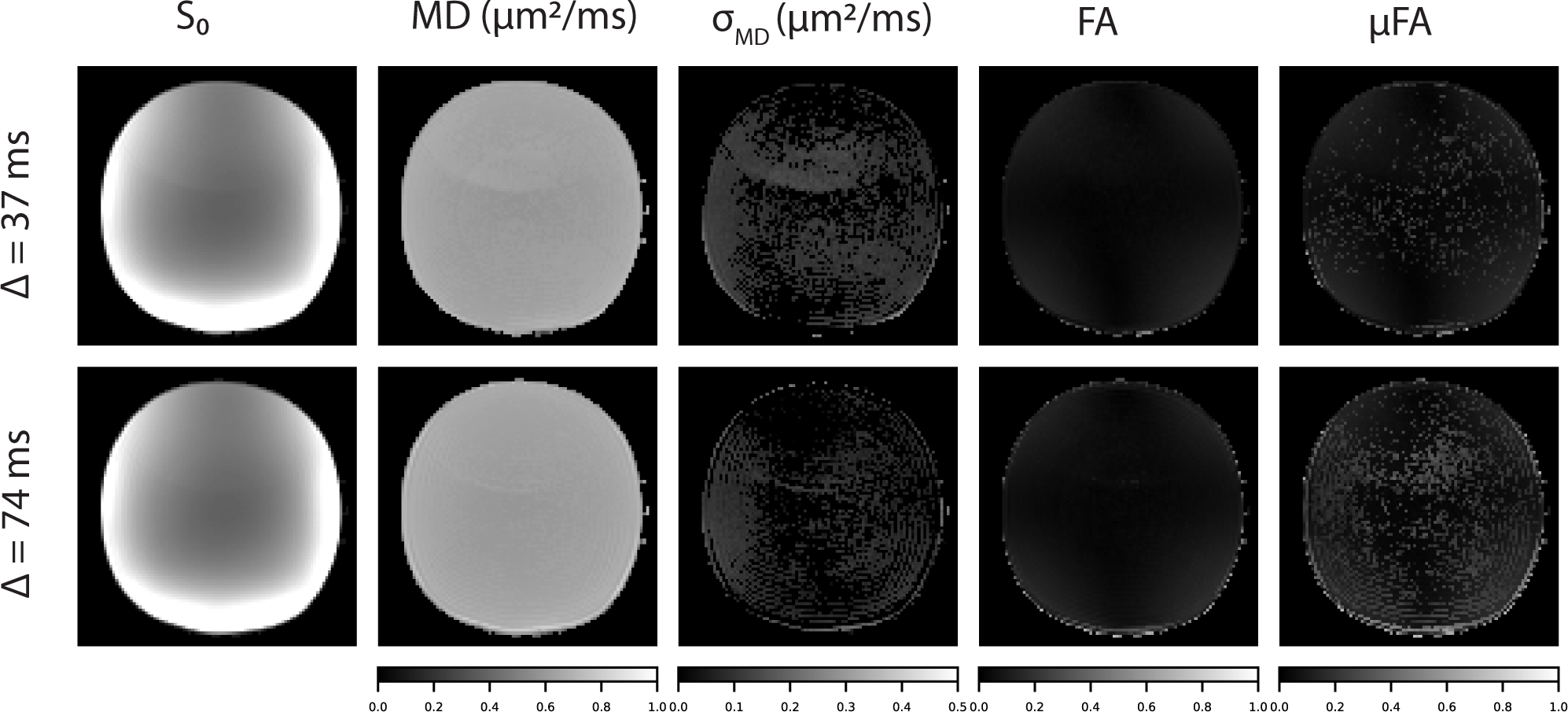
Microstructural measures derived from DTD MRI on an axial slice in the PVP phantom acquired at two different diffusion times (i.e., Δ = 37*\*74 ms). The measures include the non-diffusion weighted signal (*S*_0_), mean diffusivity (MD), standard deviation of mean diffusivity (*σ*_MD_), fractional anisotropy (FA) and microscopic fractional anisotropy (*µ*FA). The measures were uniform across the field of view except for small amounts of ghosting and gradient hardware artifacts at the boundaries with their values matching with those expected.

**Table 1:** Comparison of DTI and DTD measures obtained at two different diffusion times in a homogeneous spherical polyvinylpyrrolidone (PVP) diffusion MRI phantom. The measures are averaged over a region of interest (ROI) at the center of the phantom. Because of the homogeneity of the diffusion properties, the true values of standard deviations if the MD, FA, and *µ*FA are zero. Nonetheless, they are affected by noise in the measurements albeit this contribution is also close to zero. The difference in MD is less than 2% across the diffusion times whose value matches the expected diffusivity of PVP at the bore temperature.

| Diffusion time | MD ($\mu m^2/ms$) | $\sigma_{MD}(\mu m^2/ms)$ | FA | μFA |
| --- | --- | --- | --- | --- |
| $\Delta = 37$ ms | 0.58 | 0.02 | 0.03 | 0.04 |
| $\Delta = 74$ ms | 0.59 | 0.01 | 0.03 | 0.04 |

The raw DWIs obtained in the human brain *in vivo* for both diffusion times along with the absolute difference images and their distribution are shown in Figure 4. The signal contrast in the brain parenchyma for all b-tensors was very similar across diffusion times as verified by the white noise appearance of the absolute difference maps except at low b-values. This includes white matter regions such as corpus callosum and internal capsule fibers where water is usually assumed to be restricted. At low b-values, the larger signal differences were noted in CSF-filled regions such as in the ventricles and subarachnoid spaces. The signal differences between diffusion times became progressively smaller with increasing b-value due to the high diffusivity of CSF. This effect is manifest as fatter tails in the distribution of absolute image intensity differences at low b-values. The DTI and DTD measures obtained in the human brain *in vivo* for both diffusion times are shown in Figure 5. Given the similar signal trends, the maps of the model measures were indistinguishable. We performed quantitative ROI analysis on the cerebral cortex, corpus callosum, and lateral ventricles representative of gray matter, white matter, and CSF regions, respectively, to verify the similarity of the model measures and report the findings in Table 2. The average values of the measures were nearly identical for gray and white matter regions, albeit with increased MD and *σ*_MD_ in the gray matter ROI, indicating slight contamination of this region with CSF. In the CSF, difference in MD, *σ*_MD_, FA and *µ*FA between the two diffusion times was approximately 2%, 2 %, 42%, and 38 %, respectively.

**Figure 4:**
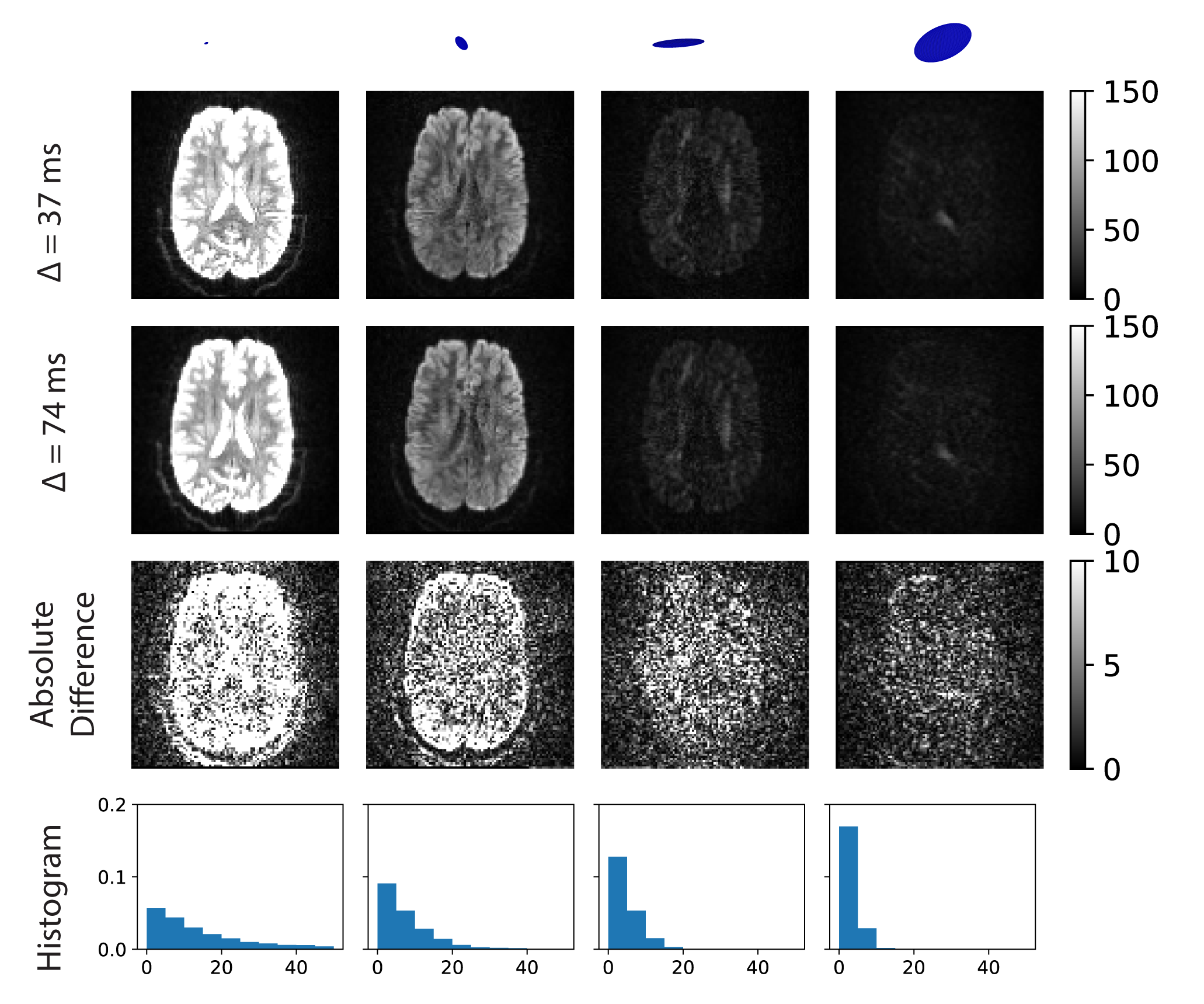
Multiple diffusion encoded (MDE) images from an axial slice in the human brain of a representative subject acquired *in vivo* at two different diffusion times (i.e., Δ = 37*\*74 ms). The rank-1/rank-2 b-tensors corresponding to each column are displayed as sticks and ellipses, respectively. The b-values of the plotted b-tensors are approximately 270, 1100, 3500, and 4820 *s/mm*^2^ respectively. The absolute difference between the two images along with the corresponding histogram are shown in the third and fourth rows respectively. The image intensities in the brain parenchyma across the diffusion times were indistinguishable as shown by the white noise like appearance of difference maps especially at large b-values. Signal variations were however observed in CSF filled ventricles likely due to pulsatile flow in these regions.

**Figure 5:**
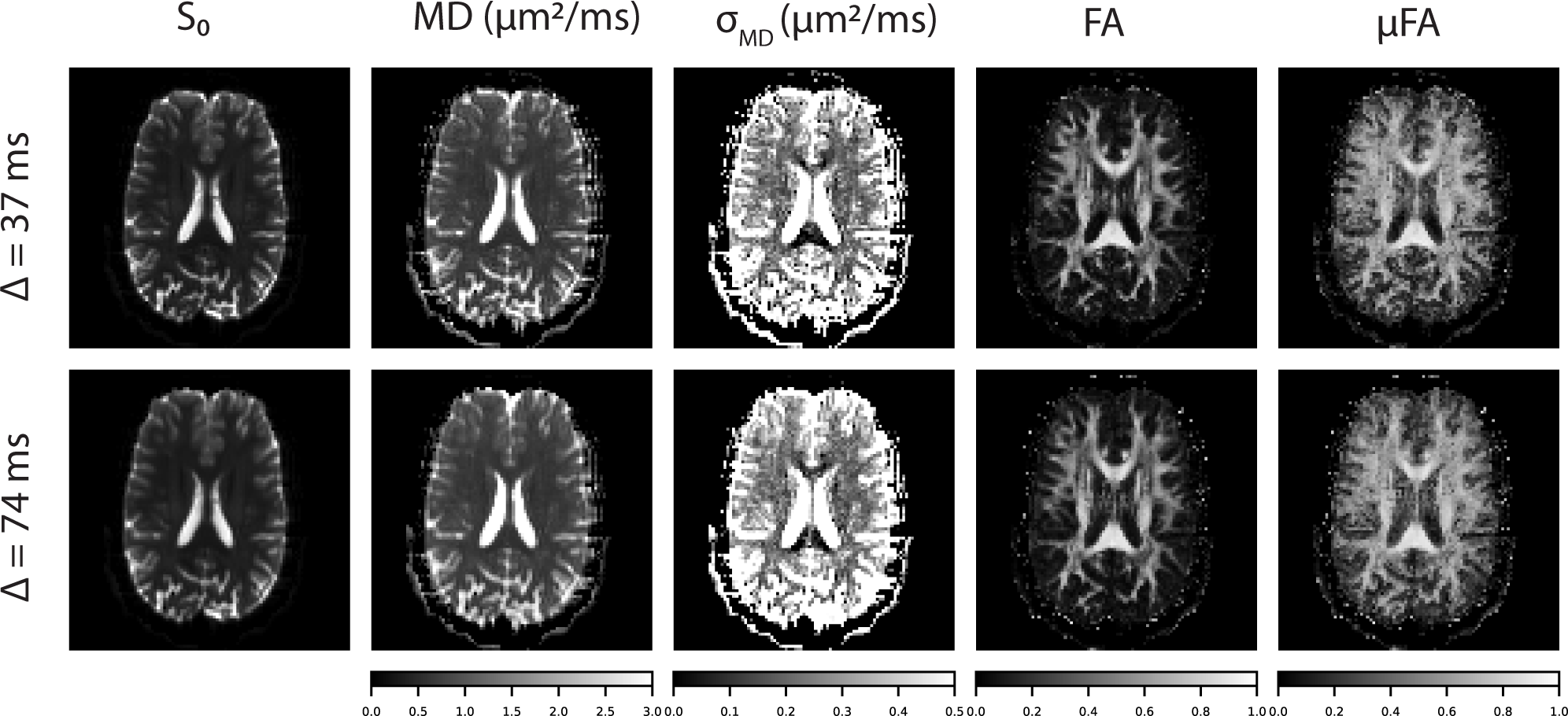
Microstructural measures derived from DTD MRI in an axial slice in the human brain acquired *in vivo* at two different diffusion times (i.e., Δ = 37*\*74 ms). The measures include the non-diffusion weighted signal (*S*_0_), mean diffusivity (MD), standard deviation of mean diffusivity (*σ*_MD_), fractional anisotropy (FA) and microscopic fractional anisotropy (*µ*FA). It can be observed that the value of the measures were as expected and nearly indistinguishable across the diffusion times tested.

**Table 2:**
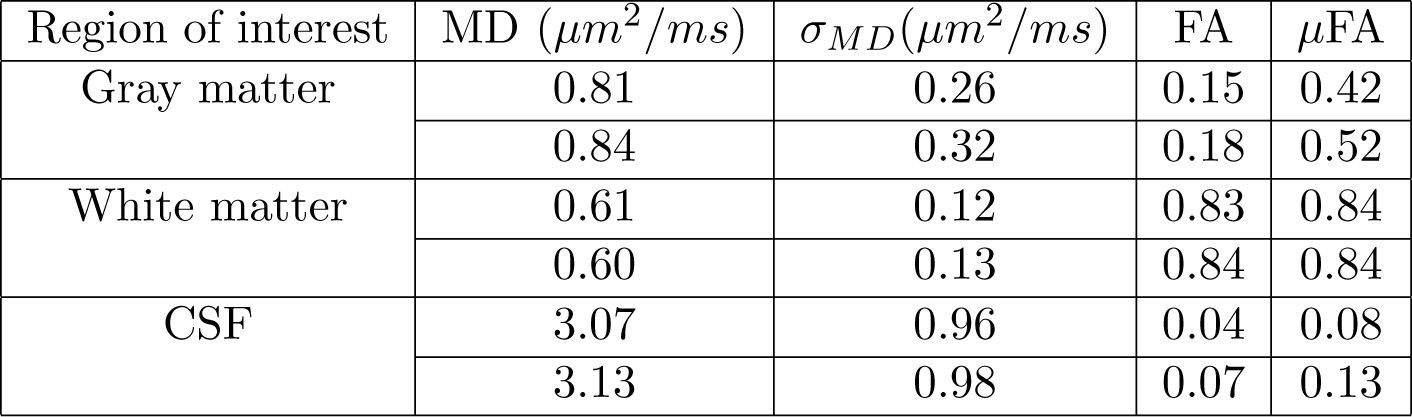
Comparison of DTI and DTD measures obtained at two different diffusion times in a brain of a representative subject. The values of the DTD and DTI measures averaged over ROIs in gray matter, white matter and CSF are reported in the above table. The two rows within each ROI represent the values at the two different diffusion times. Gray matter ROI encompasses a small region of the cerebral cortex, white matter ROI was drawn over a portion of the corpus callosum, and CSF ROI was drawn over one of the lateral ventricles. The differences in the values of these measures in gray and white matter are negligible. The diffusivity in the CSF due to the pulsation and low signal-to-noise at large b-values exhibited more variability.

| Region of interest | MD ($\mu m^2/ms$) | $\sigma_{MD}(\mu m^2/ms)$ | FA | μFA |
| --- | --- | --- | --- | --- |
| Gray matter | 0.81 | 0.26 | 0.15 | 0.42 |
|  | 0.84 | 0.32 | 0.18 | 0.52 |
| White matter | 0.61 | 0.12 | 0.83 | 0.84 |
|  | 0.60 | 0.13 | 0.84 | 0.84 |
| CSF | 3.07 | 0.96 | 0.04 | 0.08 |
|  | 3.13 | 0.98 | 0.07 | 0.13 |

The applicability of our findings across subjects is verified by mapping the absolute difference in the raw DWIs obtained at the two diffusion times for all the subjects scanned at a similar location in their brains, as shown in Figure 6 along with the *S*_0_ maps. The signal differences were observed in CSF-filled regions at low b-values, as described previously. The differences observed in the parenchyma were an order of magnitude lower than the *S*_0_ and had a white noise appearance.

**Figure 6:**
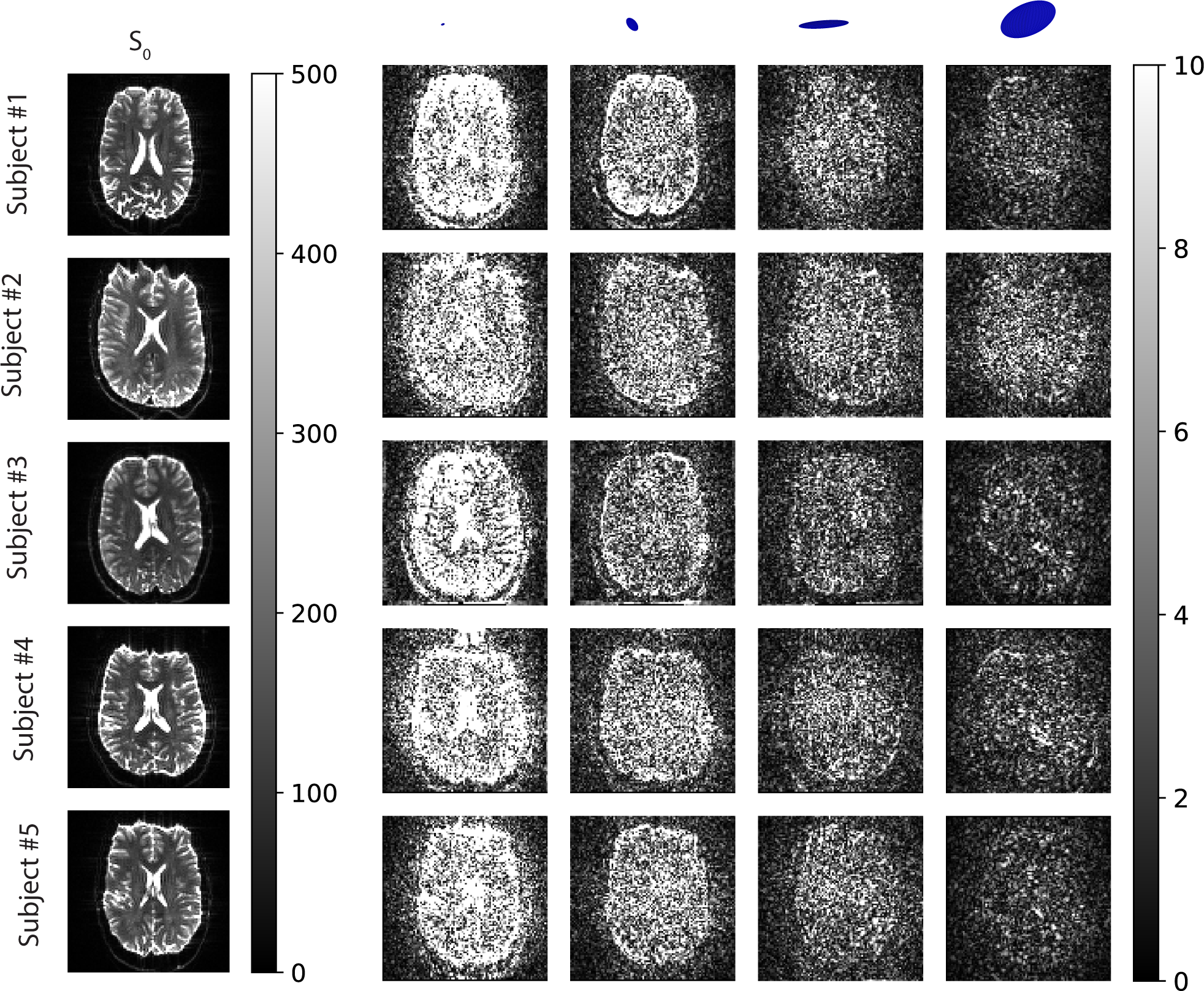
Absolute difference in multiple diffusion encoded (MDE) images relative to *S*_0_ from an axial slice in the human brain of five healthy controls *in vivo* acquired at two different diffusion times (i.e., Δ = 37*\*74 ms). The rank-1/rank-2 b-tensors corresponding to each column are displayed as sticks and ellipses, respectively. The b-values of the plotted b-tensors are approximately 270, 1100, 3500, and 4820 *s/mm*^2^ respectively. The non-diffusion weighted image (*S*_0_) for each subject is provided for reference. The DWIs acquired with two diffusion times were nearly indistinguishable except at the lateral ventricles for all the subjects scanned which further strengthen the validity of our conclusion.

## 4 Discussion

In this study, we investigate the nature of diffusion of water molecules in living human brain tissue at the mesoscopic length and time scales to arrive at a comprehensive model for the diffusion-weighted MR signal that can in turn be used to infer mesoscopic brain architectural features *in vivo*. We chose to acquire data on the 3T Connectome MRI scanner because its 300 mT/m gradients enable measurements with sensitivity to smaller diffusion displacements and higher spatial resolution at shorter echo times than would be achievable on a conventional clinical MRI scanner. The hypothesis we tested in this study is whether the Gaussian diffusion assumption is justified at the mesoscopic length scale. This was ascertained by varying a single scanning parameter (i.e. the diffusion time) while fixing the others. It is important to note that because the interfused diffusion MRI pulse sequence has a well-defined diffusion time and gradient pulse width, we were able to vary the diffusion time systematically in this study. This, for instance, would not be possible using an MRI experiment employing “free waveform” DWI acquisitions [28, 39], as these MRI pulse sequences do not have well-defined diffusion times or pulsed gradient durations that can be systematically varied. Our pulse sequence also allowed us to access shorter time scales at high SNR than those possible with the traditional mPFG sequences involving double spin-echo preparation [40].

### 4.1 Scanner quality assurance

We chose to use a PVP phantom with well-characterized diffusion properties [34] to assess the data quality acquired on the Connectome scanner. The high SNR available in the PVP phantom, whose mean diffusivity is matched to that of brain tissue, increases the visibility of artifacts often not seen *in-vivo*, thereby establishing the limits of the scanner capabilities. An idealized scanner would produce a homogeneous *S*_0_ map and mean diffusivity equal to that expected for the chosen PVP concentration and ambient bore temperature. The remaining DTI and DTD parameters would be equal to zero since there is no macroscopic or microscopic heterogeneity in the phantom. The PVP data we collected was not too far from ideal with slight imperfections that were not observed *in-vivo*. The values of the mean diffusivity and FA matched the expected values, while slight artifacts were visible in the DTD measures which were sensitive to high b-value data. The increased signal difference between the diffusion times at low b-values could be due to coherence artifacts resulting from the long relaxation times of liquids compared to brain tissue. The heterogeneity observed in the DTI and DTD maps at the edges of the phantom could be due to the nonlinearity of the strong gradients, which causes the b-tensor to deviate from the prescribed values, especially at large b-values. The ghosting observed in the maps is likely due to parallel imaging/EPI artifacts and eddy currents from diffusion gradients interacting with the imaging gradients. It should be noted that these effects were small and not observed *in-vivo*.

### 4.2 Quest for a comprehensive model

Given the reliability of the DWI data collected on the scanner, we discuss the need for a comprehensive model of water diffusion in live brain tissue pertinent to the spatiotemporal scales probed in the measurement. While brain tissue is organized over myriads of length and time scales [41], the measurement parameters typically limit the sensitivity of diffusion MRI ranging from a few microns to mm in length scale and tens of ms to one second in time scale. The three different length scales that define the regime that the PFG-NMR signal probes are the diffusion length, which indicates the distance the spins travel during the gradient (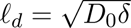), the structural length, which is length of the feature probed (*ℓ_s_*), and the gradient dephasing length, which indicates the distance spins_1_ have to travel under the gradient pulse to get dephased (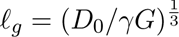) [42]. A successful model for the measured signal should be the most parsimonious one that adequately describes the microstructure at the sensitized spatiotemporal scales over a wide range of voxel motifs (*i.e.*, Occam’s razor).

For the measurement parameters used in this study, the diffusion and dephasing lengths are both approximately 3 *µ*m. The structural length (i.e., the diameter) of the constituents of central nervous system (CNS) tissue include axons (*≈* 0.1 *−* 10*µ*m with a majority less than 1*µ*m [43]), neuronal soma (3 *−* 20*µ*m), dendrites (less than 1*µ*m), astrocytes (10*µ*m soma and sub-micron processes [44]), and microglia (10*µ*m [45]). It is clear that the structural length scales in the brain are much smaller or on the same order as the diffusion and dephasing length scales used in typical clinical dMRI studies. In this so-called motional-narrowing regime, the diffusion gradients do not produce sufficient signal dephasing within each pore, but the central limit theorem results in a Gaussian phase distribution of the MR signal on an ensemble consisting of approximately 55.5 *×* 10*^−^*^6^ *×* Avogadro’s number of water molecules per cubic mm. The exact signal behavior for a Gaussian distributed phase ensemble would depend on the water permeability of the neural tissue. A highly permeable tissue would result in a Gaussian diffusion model, while impermeable boundaries lend themselves to Neumann-type solutions [46].

### 4.3 Exchange

Exchange across tissue boundaries is a vital part of any living system in order to maintain homeostasis, and as one of the most metabolically active organs in the body, the brain is no exception. Over the years, several mechanisms have been identified that facilitate water exchange in the brain [47]. Primary among these are the aquaporin (AQP) channels that are expressed on the feet of astrocytes, one of the most abundant cell types in the brain [48]. The other major water co-transporter is the Na^+^-K^+^ pump expressed in both glia and neurons that transports water while shuttling the sodium and potassium ions [49, 50].

The permeability of the tissue observed in a dMRI experiment depends on the length and time scales probed, as the exchange rate depends on the surface-to-volume ratio of the structure. In the very “short” diffusion time regime (Δ *<* 10 ms), there is not enough time for the water molecules to exchange with neighboring water pools, and the measured diffusivity approaches that of the free water [51]. As the diffusion time increases, the water molecules encounter barriers and exchange with their surroundings, thereby lowering their apparent diffusivity. This effect is observed in the oscillating gradient spin echo (OGSE) measurements, which allows probing shorter time scales than PGSE, performed on the human brain *in vivo* [16]. It should however be noted that OGSE studies are often associated with coarser length scales (i.e., low b-value) since it is difficult to probe both short length and time scales in practice due to the limitations of the gradient hardware. At very “long” diffusion times, continual exchange processes will expand the microvoxel size towards the nominal voxel size, resulting in a homogeneous voxel with a single diffusivity (i.e., *p*(D) = *δ*(D−D̅)). This regime is also difficult to probe given the finite relaxation times of water in brain tissue and reduced SNR at longer diffusion times.

In this study, we probed the intermediate length and time scales commonly used in dMRI studies because of the challenges of probing the extremes. At these scales, the brain tissue may be regarded as permeable given the multitude of cellular processes encompassed within a micro-voxel. New gradient hardware developments such as embodied in the Connectome 2.0 [52] should allow more robust probing of spatiotemporal scales. This would require a modification to the DTD model to include exchange, which is a current subject of our research. As the exchange rate increases with microvoxel size, it should be noted that there could be a point where the reduced microvoxel size afforded by strong gradients can no longer keep up with the increased exchange rate.

### 4.4 Gaussian diffusion at the mesoscale

Given the motional narrowing regime of the MR signal and permeability of the tissue at the probed length and time scales (*i.e.*, the mesoscale), an appropriate model to explain the measured dMRI signal is that of Gaussian diffusion. This is indeed evidenced by the success of DTI, which assumes Gaussian diffusion at the macroscopic scale with applications ranging from diagnosing stroke [53] and tumors [54] to neuropsychiatric conditions such as schizophrenia [55]. Despite its generality, DTI fails to elucidate the underlying microstructure in certain regions of the brain such as those consisting of crossing white matter fibers. Additionally, the measured signal in brain tissue deviates from the DTI model at sufficiently strong diffusion weightings, suggesting the presence of structural heterogeneity and/or restricted diffusion. This has led to several different models proposed to address these deficiencies with varying levels of complexity and success, including some that assume non-Gaussian diffusion [12, 13, 56].

Given the success of DTI applied to macroscopic voxels, several groups have extended DTI to the mesoscopic length scale, which has led to the DTD model [28, 30, 31]. Using novel diffusion encoding (*i.e.*, planar, spherical, etc.), DTD MRI was shown to address deficiencies of the DTI model by distinguishing microscopic and macroscopic anisotropy and explaining the diffusion-weighted signal at large b-values [28, 29, 57]. Surprisingly, a critical requirement of using the DTD framework, which is the Gaussian diffusion assumption at the mesoscopic length scale, was never addressed (*i.e.*, validated) until now. Specifically, a violation of this assumption would invalidate the use of the exponential kernel in the DTD MRI signal model (*i.e.*, Equations (1) and (2)) [28, 30].

A hallmark of Gaussian diffusion is the time independence of apparent diffusivity. A purely restricted diffusion process thought to occur in white matter fibers [58, 59] would result in an asymptotic decay in the apparent diffusivity to zero as a function of diffusion time. Our results at large enough b-values with multiple diffusion encoding show the time independence of the measured MR signal over the two diffusion times tested both in gray and white matter. Not surprisingly both the DTD and DTI measures also do not vary significantly when doubling the diffusion time.

Our results confirm previous findings on the time independence of DWI [60] and DTI measures in the human brain *in vivo* [15, 61] while extending this assumption into the mesoscopic DTD regime. It is quite remarkable that the voxel-averaged MD is not diffusion time-dependent over this range of diffusion times and thus satisfies the Einstein equation. More remarkable is that *p*(D) appears to be diffusion time independent over this range. This means that the covariance tensor, which reflects microscopic heterogeneity within the voxel, is constant over this range of diffusion times as well. This adds support to the notion that the individual micro-voxels also exhibit Gaussian diffusion in this diffusion time regime, not just their ensemble-averaged diffusivity.

It should be noted that many studies reporting time-dependent diffusion in the brain are conducted on excised brain tissue specimens [62, 63, 64] whose microstructure and water permeability are significantly altered compared to live tissue due to fixation processes [65] *inter alia*. Studies that do report time-dependent diffusion in live human white matter show very minimal effect over an even wider range of diffusion times [14].

### 4.5 Free vs. Hindered Diffusion

Classically, two types of diffusion processes exhibit Gaussian diffusion behavior over a large range of diffusion times: free and hindered diffusion. The former describes the diffusion of a liquid or gas in which there are no fixed boundaries or obstructions present to alter the displacement history of individual molecules. The latter describes diffusion in tortuous media in which there may be permeable barriers or boundaries. These two processes both follow the Einstein equation at “long” diffusion times but may exhibit different mean-squared net displacement vs. diffusion time behavior at “short” diffusion times. In this experiment, we were not able to probe this short diffusion time regime, which would enable us to distinguish between these two mechanisms of diffusion in the brain parenchyma. Because the free diffusivity of water is substantially higher than the MD we measured in tissue, we infer that we are observing hindered diffusion in the brain parenchyma.

One might expect that we would measure free diffusion in CSF compartments, such as those surrounding the cortex or within ventricles, but this was not the case. The ventricles and subarachnoid spaces are not static structures containing mostly ionized water. Pulsation artifacts in the blood and CSF compartments and periodic respiratory effects result in time-dependent advection processes to which the spin echo sequence is also susceptible, resulting in deviations from Gaussian diffusion [66]. In fact, when we first developed DTI, we found that the greatest variance of the tensor-derived eigenvalues was found in the CSF compartments of all brain tissues [11]. It was only by normalizing the standard deviation of the eigenvalues by the MD [67] that the apparently high diffusion anisotropy of the CSF was suppressed. This non-Gaussian diffusion behavior of CSF suggests that simple “free water elimination” strategies [68] would not be successful in the ventricles, although such approaches may be better suited for removing CSF signals in the brain parenchyma itself.

### 4.6 Beyond the mesoscale

A natural consequence of our findings is to ask over what range of finer length scales the Gaussian diffusion assumption remains valid. Specifically, is it valid at the micro and nano scales approaching the resolution of electron microscopy, *in vivo*? The microvoxel size required for the central limit theorem and thus the Gaussian diffusion assumption to break down is around 1 nm on each edge, which would roughly contain about 30-40 water molecules. This limit would never be achieved in practice as it would require gargantuan gradient strengths while fast exchange that occurs at these length scales would inevitably increase the actual size of the microvoxels [33]. However, between the mesoscale and nanoscale, the Gaussian diffusion assumption is expected to hold and may reveal previously invisible microstructural details, especially with the advent of next-generation diffusion MRI scanners [52].

## 5 Conclusion

Our *in vivo* human brain experiments revealed no significant measurable differences between the MDE or mPFG signal attenuation over a practical range of diffusion times (37 −74 ms) typically used in clinical imaging. These results support the argument that over this range of diffusion times in living brain tissue, the Gaussian diffusion assumption/approximation holds for multiple mesoscopic water pools simultaneously. Our findings imply that diffusion in tissues can be modeled using a mixture of Gaussian processes, which is a fundamental assumption of DTD-MRI. Thus, our findings justify the use of DTD-MRI for clinical applications over the broad range of diffusion times we investigated.

## 6 Acknowledgments

We would like to thank Teddy Cai for helpful discussions. This study was supported by the Intramural Research Program of the NICHD. This work was also partly funded by NIH BRAIN Initiative grant 1U01EB026996-01 -”Connectome 2.0: Developing the next generation human MRI scanner for bridging studies of the micro-, meso- and macro-connectome.” and by the Department of Defense in the Military Traumatic Brain Injury Initiative under award, HU0001-22-2-0058. This work utilized computational resources of the NIH HPC Biowulf cluster (http://hpc.nih.gov). The authors have no conflicts of interest to disclose. The views, information or content, and conclusions presented do not necessarily represent the official position or policy of, nor should any official endorsement be inferred on the part of, the Uniformed Services University, the Department of Defense, the U.S. Government or the Henry M. Jackson Foundation for the Advancement of Military Medicine, Inc.

## Author Contributions

KNM and PJB designed the research study and drafted the manuscript. KNM wrote the pulse sequence and performed the DTD analysis. KNM, AVA, and PJB interpreted the results. SYH acquired the MRI data. TW helped with the pulse sequence. All authors reviewed and edited the manuscript.

## References

[1] Axelrod D., Koppel D. E., Schlessinger J., Elson E., Webb W. W.. Mobility measurement by analysis of fluorescence photobleaching recovery kinetics. Biophysical Journal. 1976;16(9):1055–1069.

[2] Saxton Michael J., Jacobson Ken. SINGLE-PARTICLE TRACKING:Applications to Membrane Dynamics. Annual Review of Biophysics and Biomolecular Structure. 1997;26(1):373– 399.

[3] Bohland Jason W., Wu Caizhi, Barbas Helen, et al. A Proposal for a Coordinated Effort for the Determination of Brainwide Neuroanatomical Connectivity in Model Organisms at a Mesoscopic Scale. PLoS Computational Biology. 2009;5(3).

[4] Kac Mark. Can One Hear the Shape of a Drum?. The American Mathematical Monthly. 1966;73(4P2):1–23.

[5] Nicholson C., Tao L.. Hindered diffusion of high molecular weight compounds in brain extracellular microenvironment measured with integrative optical imaging.. Biophysical journal. 1993;65(6):2277–90.

[6] Leuze Christoph, Aswendt Markus, Ferenczi Emily, et al. The separate effects of lipids and proteins on brain MRI contrast revealed through tissue clearing. NeuroImage. 2017;156:412– 422.

[7] Williamson Nathan H., Ravin Rea, Benjamini Dan, et al. Magnetic resonance measurements of cellular and sub-cellular membrane structures in live and fixed neural tissue. eLife. 2019;8.

[8] Moseley M. E., Cohen Y., Mintorovitch J., et al. Early detection of regional cerebral ischemia in cats: Comparison of diffusion- and T2-weighted MRI and spectroscopy. Magnetic Resonance in Medicine. 1990;14(2):330–346.

[9] Basser Peter J., Mattiello James, LeBihan Denis. Estimation of the Effective Self-Diffusion Tensor from the NMR Spin Echo. Journal of Magnetic Resonance, Series B. 1994;103(3):247– 254.

[10] Basser P.J., Mattiello J., LeBihan D.. MR diffusion tensor spectroscopy and imaging. Biophysical Journal. 1994;66(1):259–267.

[11] Pierpaoli C, Jezzard P, Basser P J, Barnett A, Di Chiro G. Diffusion tensor MR imaging of the human brain.. Radiology. 1996;201(3):637–648.

[12] Jensen Jens H., Helpern Joseph A., Ramani Anita, Lu Hanzhang, Kaczynski Kyle. Diffusional kurtosis imaging: The quantification of non-gaussian water diffusion by means of magnetic resonance imaging. Magnetic Resonance in Medicine. 2005;53(6):1432–1440.

[13] Hall Matt G., Barrick Thomas R.. From diffusion-weighted MRI to anomalous diffusion imaging. Magnetic Resonance in Medicine. 2008;59(3):447–455.

[14] Fieremans Els, Burcaw Lauren M., Lee Hong Hsi, Lemberskiy Gregory, Veraart Jelle, Novikov Dmitry S.. In vivo observation and biophysical interpretation of time-dependent diffusion in human white matter. NeuroImage. 2016;129:414–427.

[15] Clark Chris A., Hedehus Maj, Moseley Michael E.. Diffusion time dependence of the apparent diffusion tensor in healthy human brain and white matter disease. Magnetic Resonance in Medicine. 2001;45(6):1126–1129.

[16] Baron Corey A., Beaulieu Christian. Oscillating gradient spin-echo (OGSE) diffusion tensor imaging of the human brain. Magnetic Resonance in Medicine. 2014;72(3):726–736.

[17] Does Mark D., Parsons Edward C., Gore John C.. Oscillating gradient measurements of water diffusion in normal and globally ischemic rat brain. Magnetic Resonance in Medicine. 2003;49(2):206–215.

[18] Moonen Chrit T.W., Pekar James, Vleeschouwer Marloes H.M., Gelderen Peter, Zijl Peter C.M., Despres Daryl. Restricted and anisotropic displacement of water in healthy cat brain and in stroke studied by NMR diffusion imaging. Magnetic Resonance in Medicine. 1991;19(2):327–332.

[19] Novikov Dmitry S., Jensen Jens H., Helpern Joseph A., Fieremans Els. Revealing mesoscopic structural universality with diffusion. Proceedings of the National Academy of Sciences. 2014;111(14):5088–5093.

[20] Lee Hong Hsi, Fieremans Els, Novikov Dmitry S.. What dominates the time dependence of diffusion transverse to axons: Intra-or extra-axonal water?. NeuroImage. 2018;182:500–510.

[21] Avram Liat, Özarslan Evren, Assaf Yaniv, Bar-Shir Amnon, Cohen Yoram, Basser Peter J.. Three-dimensional water diffusion in impermeable cylindrical tubes: theory versus experiments. NMR in Biomedicine. 2008;21(8):888–898.

[22] Komlosh Michal E., Özarslan Evren, Lizak Martin J., et al. Pore diameter mapping using double pulsed-field gradient MRI and its validation using a novel glass capillary array phantom. Journal of Magnetic Resonance. 2011;208(1):128–135.

[23] McNab Jennifer A., Edlow Brian L., Witzel Thomas, et al. The Human Connectome Project and beyond: Initial applications of 300mT/m gradients. NeuroImage. 2013;80:234–245.

[24] Basser Peter J. Fiber-Tractography via Diffusion Tensor MRI (DT-MRI). In: Annual Meeting of International Society for Magnetic Resonance in Medicine:1226; 1998; Sydney, Australia.

[25] Inglis B.A., Bossart E.L., Buckley D.L., Wirth E.D., Mareci T.H.. Visualization of neural tissue water compartments using biexponential diffusion tensor MRI. Magnetic Resonance in Medicine. 2001;45(4):580–587.

[26] Özarslan Evren, Mareci Thomas H.. Generalized diffusion tensor imaging and analytical re-lationships between diffusion tensor imaging and high angular resolution diffusion imaging. Magnetic Resonance in Medicine. 2003;50(5):955–965.

[27] Cory DG, Garroway Allen N., Miller Joel B.. Applications of spin transport as a probe of local geometry. In: Proceedings of the American Chemical Society:149–150; 1990.

[28] Westin Carl-Fredrik, Knutsson Hans, Pasternak Ofer, et al. Q-space trajectory imaging for multidimensional diffusion MRI of the human brain. NeuroImage. 2016;135:345–362.

[29] Magdoom Kulam Najmudeen, Pajevic Sinisa, Dario Gasbarra, Basser Peter J. A new framework for MR diffusion tensor distribution. Scientific Reports. 2021;11(1):2766.

[30] Jian Bing, Vemuri Baba C., Özarslan Evren, Carney Paul R., Mareci Thomas H.. A novel tensor distribution model for the diffusion-weighted MR signal. NeuroImage. 2007;37(1):164– 176.

[31] Basser P.J., Pajevic S.. A normal distribution for tensor-valued random variables: applications to diffusion tensor MRI. IEEE Transactions on Medical Imaging. 2003;22(7):785–794.

[32] Nilsson Markus, Szczepankiewicz Filip, Brabec Jan, et al. Tensor-valued diffusion MRI in under 3 minutes: an initial survey of microscopic anisotropy and tissue heterogeneity in intracranial tumors. Magnetic Resonance in Medicine. 2020;83(2):608–620.

[33] Magdoom Kulam Najmudeen, Avram Alexandru V., Sarlls Joelle E., Dario Gasbarra, Basser Peter J.. A novel framework for in-vivo diffusion tensor distribution MRI of the human brain. NeuroImage. 2023;271:120003.

[34] Pierpaoli C, Sarlls J, Nevo U, Basser Peter J., Horkay F. Polyvinylpyrrolidone (PVP) water solutions as isotropic phantoms for diffusion MRI studies. In: Annual Meeting of International Society for Magnetic Resonance in Medicine:1414; 2009.

[35] Setsompop Kawin, Gagoski Borjan A., Polimeni Jonathan R., Witzel Thomas, Wedeen Van J., Wald Lawrence L.. Blipped-controlled aliasing in parallel imaging for simultaneous multi-slice echo planar imaging with reduced g-factor penalty. Magnetic Resonance in Medicine. 2012;67(5):1210–1224.

[36] Jenkinson Mark, Smith Stephen. A global optimisation method for robust affine registration of brain images. Medical Image Analysis. 2001;5(2):143–156.

[37] Jenkinson Mark, Bannister Peter, Brady Michael, Smith Stephen. Improved Optimization for the Robust and Accurate Linear Registration and Motion Correction of Brain Images. NeuroImage. 2002;17(2):825–841.

[38] Smith Stephen M., Jenkinson Mark, Woolrich Mark W., et al. Advances in functional and structural MR image analysis and implementation as FSL. NeuroImage. 2004;23(SUPPL. 1):S208–S219.

[39] Sjölund Jens, Szczepankiewicz Filip, Nilsson Markus, Topgaard Daniel, Westin Carl-Fredrik, Knutsson Hans. Constrained optimization of gradient waveforms for generalized diffusion encoding. Journal of Magnetic Resonance. 2015;261:157–168.

[40] Mitra Partha P.. Multiple wave-vector extensions of the NMR pulsed-field-gradient spin-echo diffusion measurement. Physical Review B. 1995;51(21):15074–15078.

[41] Lichtman Jeff W., Denk Winfried. The big and the small: Challenges of imaging the brain’s circuits. Science. 2011;334(6056):618–623.

[42] Grebenkov Denis S.. NMR survey of reflected Brownian motion. Reviews of Modern Physics. 2007;79(3):1077–1137.

[43] Perge János A., Niven Jeremy E., Mugnaini Enrico, Balasubramanian Vijay, Sterling Peter. Why Do Axons Differ in Caliber?. Journal of Neuroscience. 2012;32(2):626–638.

[44] Oberheim Nancy Ann, Takano Takahiro, Han Xiaoning, et al. Uniquely Hominid Features of Adult Human Astrocytes. Journal of Neuroscience. 2009;29(10):3276–3287.

[45] Torres-Platas Susana G., Comeau Samuel, Rachalski Adeline, et al. Morphometric characterization of microglial phenotypes in human cerebral cortex. Journal of Neuroinflammation. 2014;11(1):1–14.

[46] Neuman C. H.. Spin echo of spins diffusing in a bounded medium. The Journal of Chemical Physics. 2003;60(11):4508.

[47] MacAulay N., Hamann S., Zeuthen T.. Water transport in the brain: Role of cotransporters. Neuroscience. 2004;129(4):1029–1042.

[48] Freeman Marc R., Rowitch David H.. Evolving concepts of gliogenesis: A look way back and ahead to the next 25 years. Neuron. 2013;80(3):613–623.

[49] Springer Charles S.. Using 1H2O MR to measure and map sodium pump activity in vivo. Journal of Magnetic Resonance. 2018;291:110–126.

[50] Bai Ruiliang, Springer Charles S., Plenz Dietmar, Basser Peter J.. Brain active transmembrane water cycling measured by MR is associated with neuronal activity. Magnetic Resonance in Medicine. 2018;.

[51] Mitra Partha P., Sen Pabitra N., Schwartz Lawrence M.. Short-time behavior of the diffusion coefficient as a geometrical probe of porous media. Physical Review B. 1993;47(14):8565.

[52] Huang Susie Y., Witzel Thomas, Keil Boris, et al. Connectome 2.0: Developing the next-generation ultra-high gradient strength human MRI scanner for bridging studies of the micro-, meso- and macro-connectome. NeuroImage. 2021;243:118530.

[53] Basser P.J., Mattiello J., Pierpaoli C., LeBihan D. MR diffusion tensor imaging of ischemic brain in vivo. In: Proceedings of 16th Annual International Conference of the IEEE Engineering in Medicine and Biology Society:566–567IEEE; 1994.

[54] Wieshmann U. C., Symms M. R., Parker G. J.M., et al. Diffusion tensor imaging demonstrates deviation of fibres in *}*normal appearing white matter adjacent to a brain tumour. Journal of Neurology, Neurosurgery, and Psychiatry. 2000;68(4):501.

[55] Voineskos Aristotle N., Lobaugh Nancy J., Bouix Sylvain, et al. Diffusion tensor tractography findings in schizophrenia across the adult lifespan. Brain. 2010;133(5):1494–1504.

[56] Tuch David S.. Q-ball imaging. Magnetic Resonance in Medicine. 2004;52(6):1358–1372.

[57] Callaghan P. T., Komlosh M. E.. Locally anisotropic motion in a macroscopically isotropic system: displacement correlations measured using double pulsed gradient spin-echo NMR. Magnetic Resonance in Chemistry. 2002;40(13):S15–S19.

[58] Assaf Yaniv, Blumenfeld-Katzir Tamar, Yovel Yossi, Basser Peter J.. Axcaliber: A method for measuring axon diameter distribution from diffusion MRI. Magnetic Resonance in Medicine. 2008;59(6):1347–1354.

[59] Assaf Yaniv, Basser Peter J.. Composite hindered and restricted model of diffusion (CHARMED) MR imaging of the human brain. NeuroImage. 2005;27(1):48–58.

[60] Le Bihan D, Turner R, Douek P. Is water diffusion restricted in human brain white matter? An echo-planar NMR imaging study. Neuroreport. 1993;4(7):887–90.

[61] Nilsson Markus, Lätt Jimmy, Nordh Emil, Wirestam Ronnie, Ståhlberg Freddy, Brockstedt Sara. On the effects of a varied diffusion time in vivo: is the diffusion in white matter restricted?. Magnetic Resonance Imaging. 2009;27(2):176–187.

[62] Lundell H., Nilsson M., Dyrby T. B., et al. Multidimensional diffusion MRI with spectrally modulated gradients reveals unprecedented microstructural detail. Scientific Reports. 2019;9(1).

[63] Ianuş Andrada, Jespersen Sune N., Serradas Duarte Teresa, Alexander Daniel C., Drobnjak Ivana, Shemesh Noam. Accurate estimation of microscopic diffusion anisotropy and its time dependence in the mouse brain. NeuroImage. 2018;183:934–949.

[64] Özarslan Evren, Basser Peter J., Shepherd Timothy M., Thelwall Peter E., Vemuri Baba C., Blackband Stephen J.. Observation of anomalous diffusion in excised tissue by characterizing the diffusion-time dependence of the MR signal. Journal of Magnetic Resonance. 2006;.

[65] Shepherd Timothy M, Thelwall Peter E, Stanisz Greg J, Blackband Stephen J.. Aldehyde fixative solutions alter the water relaxation and diffusion properties of nervous tissue. Magnetic Resonance in Medicine. 2009;62(1):26–34.

[66] Chen Liyong, Beckett Alexander, Verma Ajay, Feinberg David A. Dynamics of respiratory and cardiac CSF motion revealed with real-time simultaneous multi-slice EPI velocity phase contrast imaging. NeuroImage. 2015;122:281–287.

[67] Basser Peter J.. New Histological and Physiological Stains Derived from Diffusion-Tensor MR Images. Annals of the New York Academy of Sciences. 1997;820(1):123–138.

[68] Pasternak Ofer, Sochen Nir, Gur Yaniv, Intrator Nathan, Assaf Yaniv. Free water elimination and mapping from diffusion MRI. Magnetic Resonance in Medicine. 2009;62(3):717–730.

